# A pathway-centric view of spatial proximity in the 3D nucleome across cell lines

**DOI:** 10.1101/027045

**Authors:** Hiren Karathia, Carl Kingsford, Michelle Girvan, Sridhar Hannenhalli

**Affiliations:** Center for Bioinformatics and Computational Biology, University of Maryland, College Park, MD; Computational Biology Department, Carnegie Mellon University, Pittsburgh, PA; Department of Physics, University of Maryland, College Park, MD

**Author notes:** Corresponding author 3104G Biomolecular Sciences Building (#296) University of Maryland, College Park, MD 20742, USA (v) (f). Authors’ emails: Hiren Karathia – Carl Kingsford – Michelle Girvan.

**Keywords:** Hi-C, Chromatin structure, Pathways, Transcriptional regulation

## Abstract

Spatial organization of the genome is critical for condition-specific gene expression. Previous studies have shown that functionally related genes tend to be spatially proximal. However, these studies have not been extended to multiple human cell types, and the extent to which context-specific spatial proximity of a pathway is related to its context-specific activity is not known. We report the first pathway-centric analyses of spatial proximity in six human cell lines. We find that spatial proximity of genes in a pathway tends to be context-specific, in a manner consistent with the pathway’s context-specific expression and function; housekeeping genes are ubiquitously proximal to each other, and cancer-related pathways such as p53 signaling are uniquely proximal in hESC. Intriguingly, we find a correlation between the spatial proximity of genes and interactions of their protein products, even after accounting for the propensity of co-pathway proteins to interact. Related pathways are also often spatially proximal to one another, and housekeeping genes tend to be proximal to several other pathways suggesting their coordinating role. Further, the spatially proximal genes in a pathway tend to be the drivers of the pathway activity and are enriched for transcription, splicing and transport functions. Overall, our analyses reveal a pathway-centric organization of the 3D nucleome whereby functionally related and interacting genes, particularly the initial drivers of pathway activity, but also genes across multiple related pathways, are in spatial proximity in a context-specific way. Our results provide further insights into the role of differential spatial organization in cell type-specific pathway activity.

## Introduction

Recent advances in Chromosome Confirmation Capture (3C) and its high throughput derivative, Hi-C, have enabled genome-wide identification of spatially proximal genomic regions [1–3]. Comparative analysis of Hi-C data across cell lines and species reveals a conserved framework of 3D architecture, represented by topologically associating domains (TADs) and further context-specific variation in distal interactions [4].

Among other things, these 3D maps of chromosomes help explain, in part, spatio-temporal regulation of gene expression by distal enhancers, aided by long-range DNA looping [5–7]. Similarly, previous studies have shown that groups of spatially clustered enhancers exhibit coactivity across cell types and this co-activity is reflected in co-expression of proximal genes, which are often functionally related [8, 9]. More specifically, genes involved in the same pathway have been shown to be spatially proximal in *Saccharomyces cerevisiae* [10, 11], *Plasmodium falciparum* [12] and *Homo sapiens* lymphoblastoid cell lines [13]. However, these previous studies have not been extended to multiple human cell lines, and it is not clear to what extent spatio-temporal activity of pathways is related to the spatial proximity of the constituent genes. More generally, the broader characterization of physically proximity of genes in the context of functional pathways is missing and could reveal organizing principles underlying spatial proximity of pathway genes as they relate to pathway activity.

In this work, we perform a comparative pathway-centric analysis of Hi-C-derived spatial proximity data in 6 ENCODE [14] cell types - *HEK293 [15], hESC [4], IMR90, BT483[16], GM06990[16]*, and *RWPE1[17]*, each with replicate data. Our analysis of two large sets of pathways – KEGG [18], and NetPath [19] reveals several properties of spatial proximity of pathway genes: We find that in general, genes in a pathway tend to be spatially proximal and this tendency is even greater for gene pairs that belong to multiple pathways. Our expression analysis shows that genes that are co-localized in nuclear space with other genes have higher expression, and this effect is especially prominent when they are proximal to a gene in the same pathway. We also found that spatial proximity of pathway genes is strongly correlated with cell type-specific pathway activity. As an expected corollary, housekeeping genes, by virtue of being ubiquitously active, exhibit ubiquitous spatial proximity. Surprisingly though, we found that the protein products of spatially proximal genes in a pathway have a significantly greater tendency to physically interact than various controls. Functional enrichment analysis suggests that spatially proximal pathway genes are enriched for specific functional classes such as transcription factor and transmembrane genes, and they occupy higher levels in the regulatory hierarchy. Finally, we look at higher-level spatial organization of functional pathways by quantifying spatial proximity for all pairs of pathways. Using this, we identify a network of spatially proximal pathways that is consistent with their functional roles.

Overall, this first comprehensive pathway-centric analysis of spatial proximity in multiple human cell lines shows a strong link between spatial proximity and context-specific gene expression and pathway activity. Our analysis also reveals surprising links between spatial proximity and interaction between the corresponding protein products. Functional analysis of proximal genes within pathways strongly suggests a regulatory hierarchical bias in physical proximity of pathway genes. Taken together, these results are consistent with a mechanism in which early regulatory components of a pathway are brought into spatial proximity in a condition specific manner.

## Results

### 1. Software pipeline for Hi-C processing - overview

Fig. 1 shows the overall pipeline that, starting from the raw reads obtained from a Hi-C experiment, produces significant pair-wise gene interactions. Details of the pipeline are provided in the Methods section, and we highlight a few pertinent features here. The pipeline allows the user to select a resolution at which significant interactions are identified. We performed our analysis at 100 kb resolution because we are interested in gene-centric interactions and 100 kb is expected to cover ∼1 gene; smaller resolution yields fewer significant interactions due to loss in power and larger resolution results in ambiguous gene-segment mapping. We have further discussed the choice of the resolution later in the discussion section. The pipeline uses the normalization step of the Homer tool [20] to control for the genomic distance-dependent features of Hi-C counts (that proximal genomic regions are more likely to interact). Using this pipeline, we processed 6 sets of pooled replicates for 6 ENCODE cell lines - HEK293, hESC, IMR90, BT483, GM06990, and RWPE1. Table 1 shows the data obtained for the 6 cell types at the default interaction p-value threshold of 0.001 and FDR <= 0.1.

**Figure 1.**
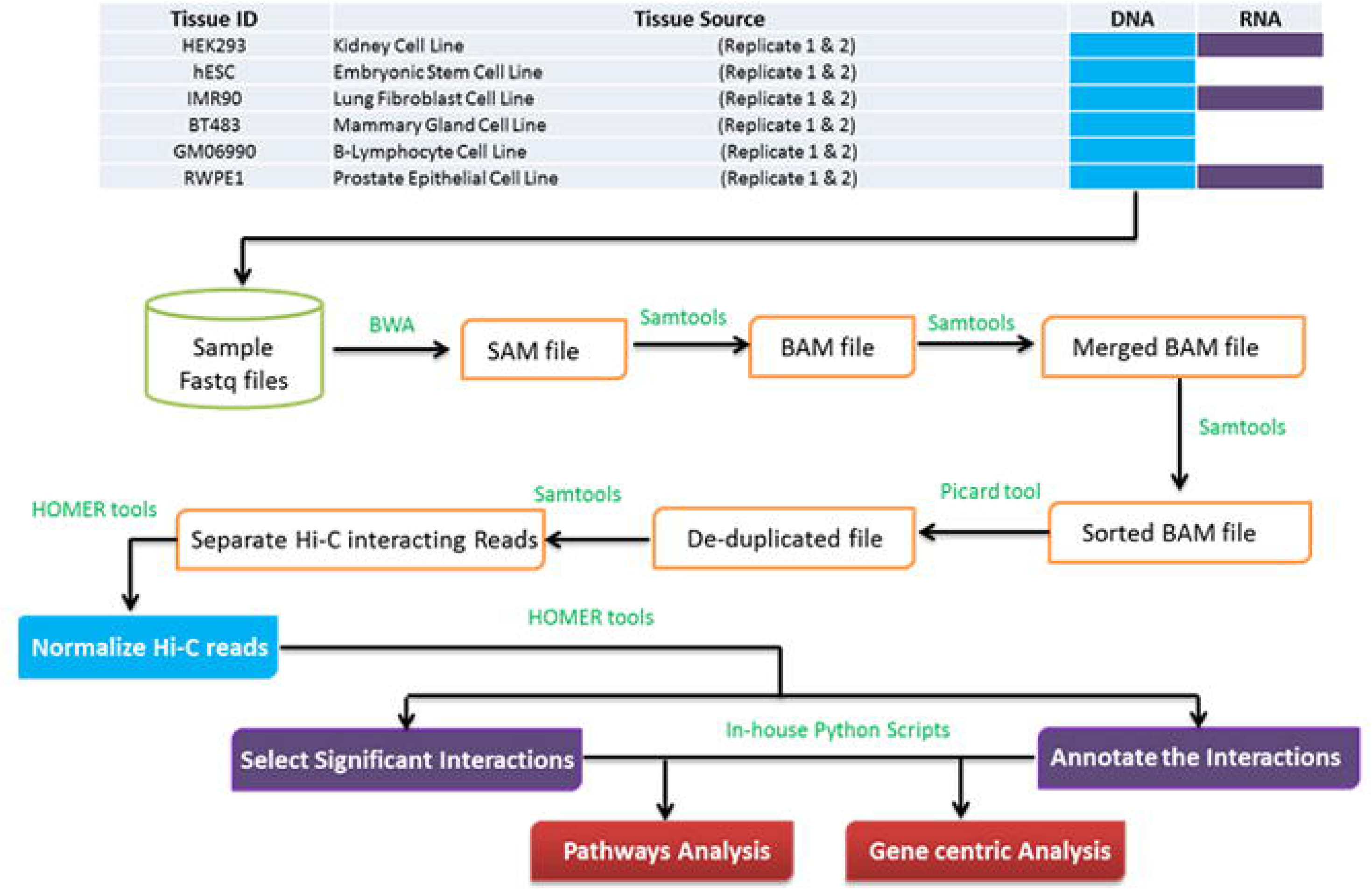
Overview of the Hi-C processing pipeline and flow of downstream analysis (see Methods for details).

**Table 1.** Hi-C analysis summary.

| Cell line | Rep-1<br>Hi-C mapped<br>fragments in<br>million | Rep-2<br>Hi-C mapped<br>fragments in<br>million | # Unique genes<br>in HiC<br>interactions | # Unique<br>genes in HiC<br>interactions |
| --- | --- | --- | --- | --- |
| IMR90 | 66.69 | 299.86 | 14550 | 101502 |
| HEK293 | 92.03 | 210 | 11978 | 44977 |
| hESC | 42.49 | 234.33 | 10713 | 34531 |
| GM06990 | 58.62 | 96.33 | 9019 | 11447 |
| BT483 | 41.8 | 64.7 | 5948 | 5792 |
| RWPE1 | 73.65 | 82.33 | 9416 | 14124 |

### Assessment of spatial proximity of pathway genes

We quantify cell-line-specific spatial proximity of genes in a pathway using *edge fraction (EF)*, which is essentially the fraction of all possible gene pairs in the pathway that are spatially proximal. This measure was previously shown to be effective [10]. We then quantify significance of EF based on a sampling procedure (see Materials & methods), obtaining a Z-score and the corresponding p-value and multiple testing corrected q-value; a higher Z-score is indicative of spatial proximity of the pathway genes above expectation. We studied 164 KEGG pathways [21] with at least 10 genes and estimated their spatial proximity Z-scores in 6 cell lines. Fig. 2 shows that overall biological pathways tend to be spatially proximal, consistent with previous reports [10]. Interestingly, as shown in Fig. 2, we found that spatial proximity for subsets of genes shared between two pathways is even greater, suggesting that such genes, coordinating multiple pathways, may be under a greater constraint to be spatial proximal.

**Figure 2.**
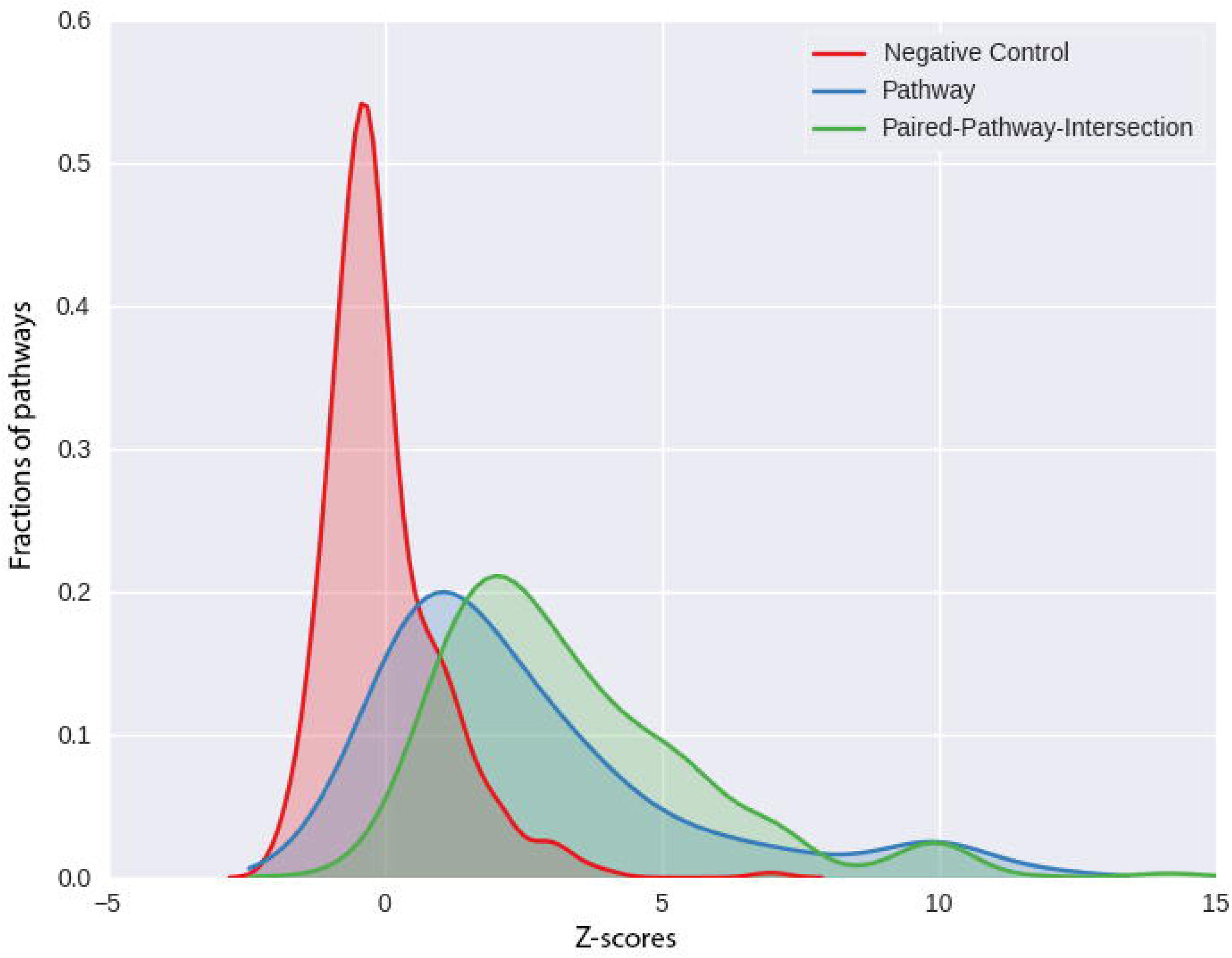
Z-score distribution of intra-pathway spatial proximity. The figure shows the distributions of spatial-proximity z-scores for three sets of gene sets, pooled from 6 cell lines. Blue: KEGG pathway. Red: Random gene-sets matching each KEGG pathway controlled for gene lengths and chromosomal distributions. Green: intersection of each pair of KEGG pathways.

Fig. 3 shows the cell-type-specific Z-scores for a representative set of pathways and Supplementary Fig. 1 shows the same for all pathways. Consistent with the fact that the KEGG database is dominated by essential and broad cellular processes, we found that spatially proximity of KEGG pathways are not only generally high (Fig. 2), but a large fraction of pathways exhibit a significant level of spatial proximity in many cell types (Fig. 4). In particular, given the ubiquitous expression and function of housekeeping genes, we tested whether these genes tend to be ubiquitously spatially proximal or whether their ubiquitous expression is decoupled from their spatial proximity to one other. Based on 3800 housekeeping genes [22], we found that housekeeping genes exhibit significant spatial proximity to other housekeeping genes in 5 out of 6 cell lines tested (Fig. 3). Other ubiquitously proximal pathways include *Metabolism of xenobiotics by cytochrome P450, 1- and 2-Methylnephthalene degradation and Gamma-hexachlorocyclohexane degradation*, all involved in drug metabolisms in animals.

**Figure 3.**
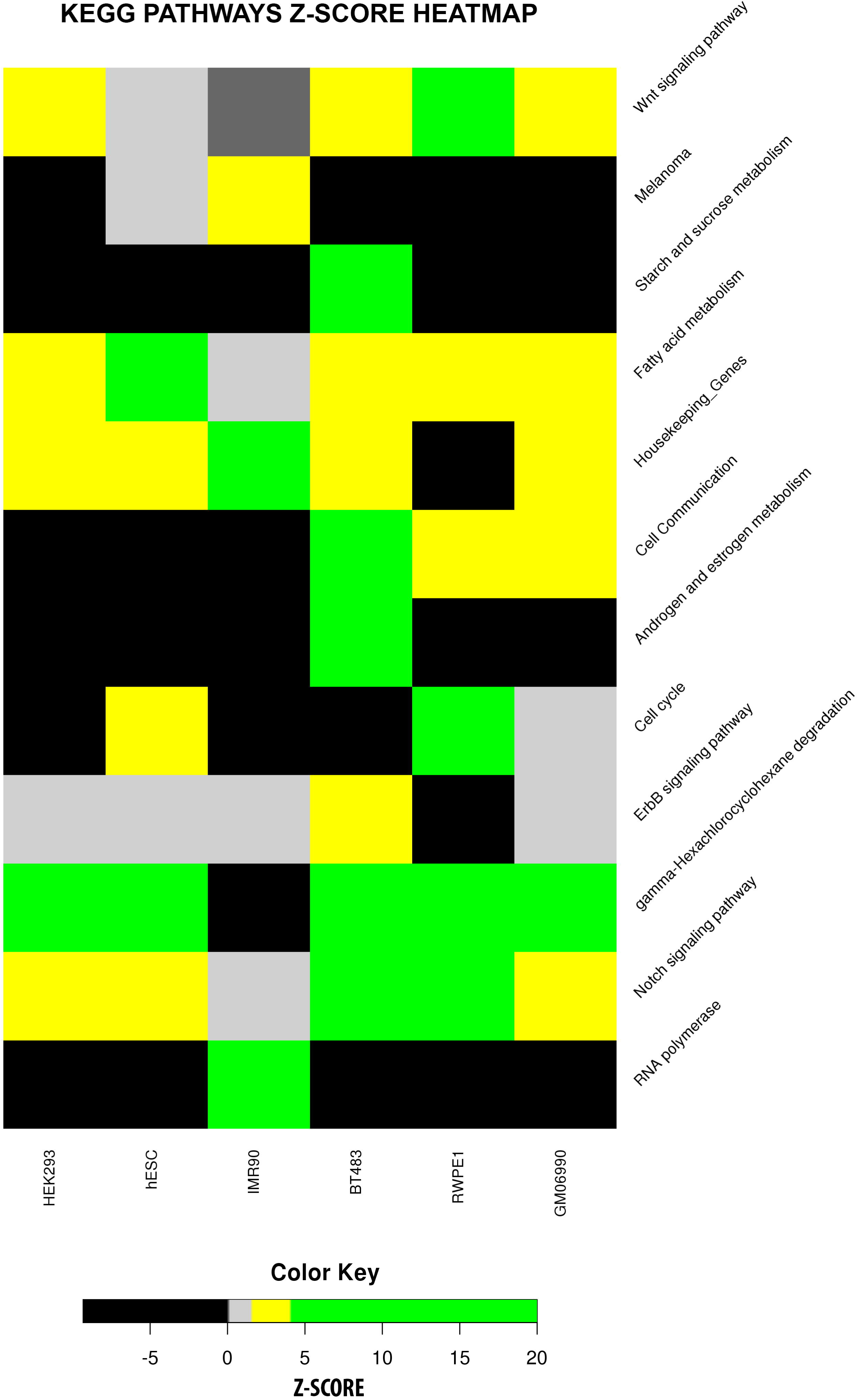
Spatial proximity of intra-pathway genes in selected KEGG pathways across six cell lines. Note the ubiquitous proximity of Housekeeping genes, hESC-specific proximity of p53 signaling pathway.

**Figure 4.**
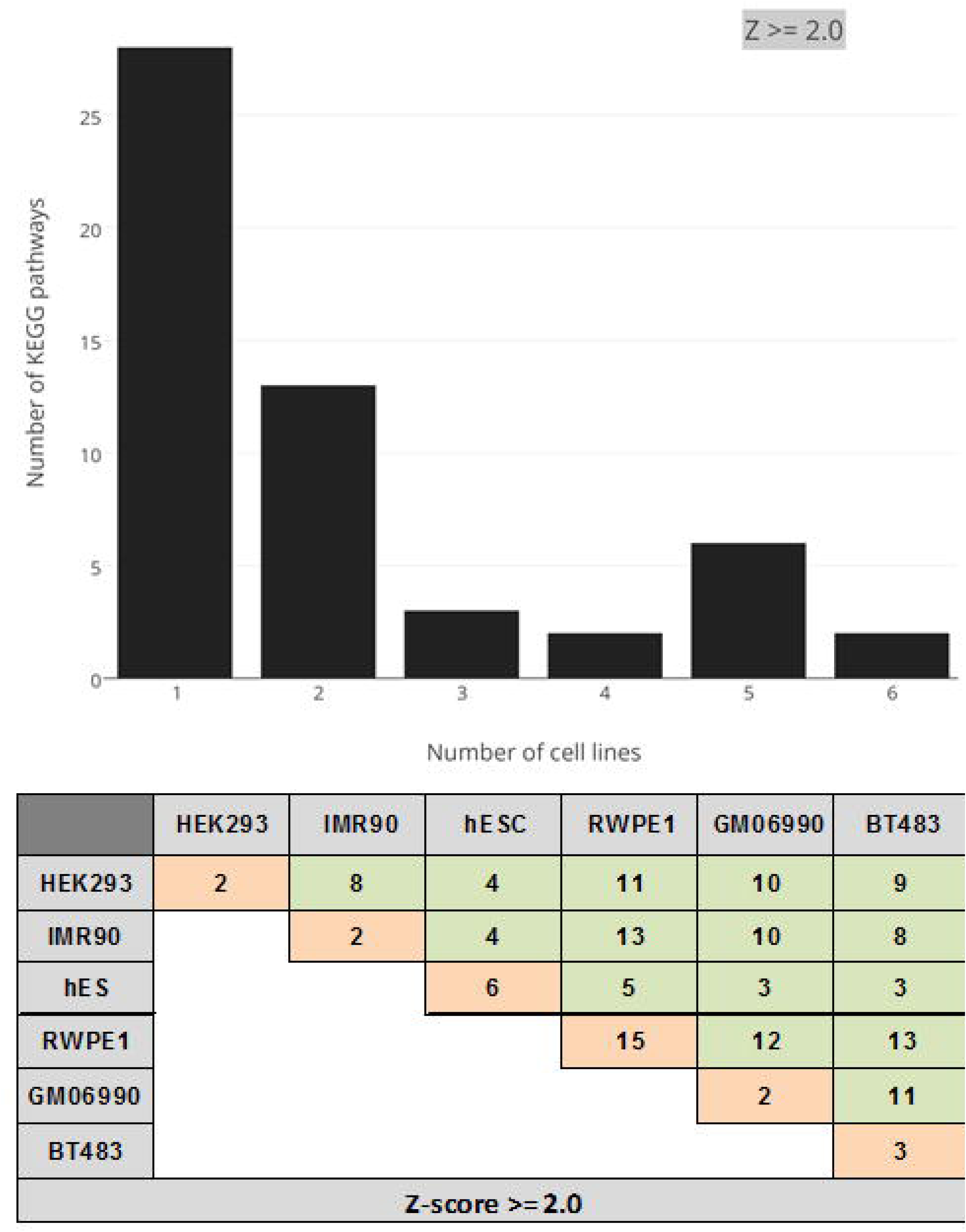
*Intra-pathway* pathway proximity is shared across tissues. The figure shows number of pathways (Y-axis) with high ***intra-pathway*** proximity (Z-score >= 2) in different number of cell lines (X-axis). The table values show number of pathways whose ***intra-pathway*** genes proximity (Z-score >= 2) is unique to a cell line (diagonal) or shared between a pair of cell lines (off-diagonal).

Several cases of cell-type-restricted spatial proximity are worth noting. For instance, ‘Cell cycle’ genes are expected to be active in pluripotent stem cells. This pathway is significantly proximal in only two cell types, one of which is the human embryonic stem cell (hESC). Cytokine-mediated signaling is critical in immune response and consistently, Cytokine-cytokine receptor interactions are uniquely proximal in immune B-cell (GM06990); B-cell-specific proximity of cytokines CCL23 and CCL4 is consistent with their known role in increased monocyte recruitment during inflammation [23]. Likewise we found the *Androgen and estrogen metabolism* pathways to be proximal in breast cancer cell lines, where the role of this pathway is well known [24]. Interestingly, we found the *Androgen-Estrogen receptor* pathway to be proximal in Kidney cell line as well (Z-score = 6.5, FDR = 0.12), consistent with the role of this pathway in glucuronidation activity that involves communication between thyroid and kidney [25]. Unexpectedly, we see proximity of ‘*Type-II diabetic mellitus*’ in lung fibroblast-derived IMR90. This is however consistent with recently observed connections between diabetes and lung functions [26]. Finally, one the only two cell lines where Melanogenesis genes is found to be proximal are the prostate epithelial RWEP1 and mammary epithelial BT483; melanogenesis in prostate epithelial cells has been previously reported [27].

As an additional set of pathways, we examined a set of annotated cancer-related pathways from NetPath [19]. As shown in Supplementary Fig. 2, genes in cancer-related pathways exhibit much more subdued spatial proximity patterns in cell lines not derived from primary tumors. Another noticeable trend is that many pathways exhibit high intra-pathway gene proximity in hESC, consistent with previously noted similarities between cancer and stem cell processes [28]. We found that *IL3-signaling* pathway is uniquely proximal in hESC consistent with its known role in cell cycle control [29].

As a useful resource, in Supplementary Table 1 we have provided, for all pathways considered, the specific gene pairs that were proximal in different cell lines.

### 2. Spatial proximity and gene expression

Next, we assessed whether spatial proximity of genes is correlated with their expression levels. Furthermore, we also assessed among the spatially proximal genes whether belonging to same pathway has any association with expression level. This analysis was done in all 6 cell lines using RNA-seq data available in GEO (see Materials & Methods). We compared cell type specific expression levels for three disjoint groups of genes (Fig. 5). The first group consisted of genes that are proximal to another gene in the same pathway (*proximal-intra-pathway*). The second group consisted of genes proximal to another gene but excluding *proximal-intra-pathway* genes; this group was designed to assess whether shared pathway membership impinges on gene expression. The last group consisted of all other genes not proximal to any gene (*non-proximal-generic*). Fig. 6 shows that in pooled result from all 6 tissues, while genes that are spatially proximal to other genes have a greater expression than non-proximal genes, the expression is greater for genes that are proximal to a gene in the same pathway. The results for individual cell lines are qualitatively similar and are shown in Supplementary Fig. 3.

**Figure 5.**
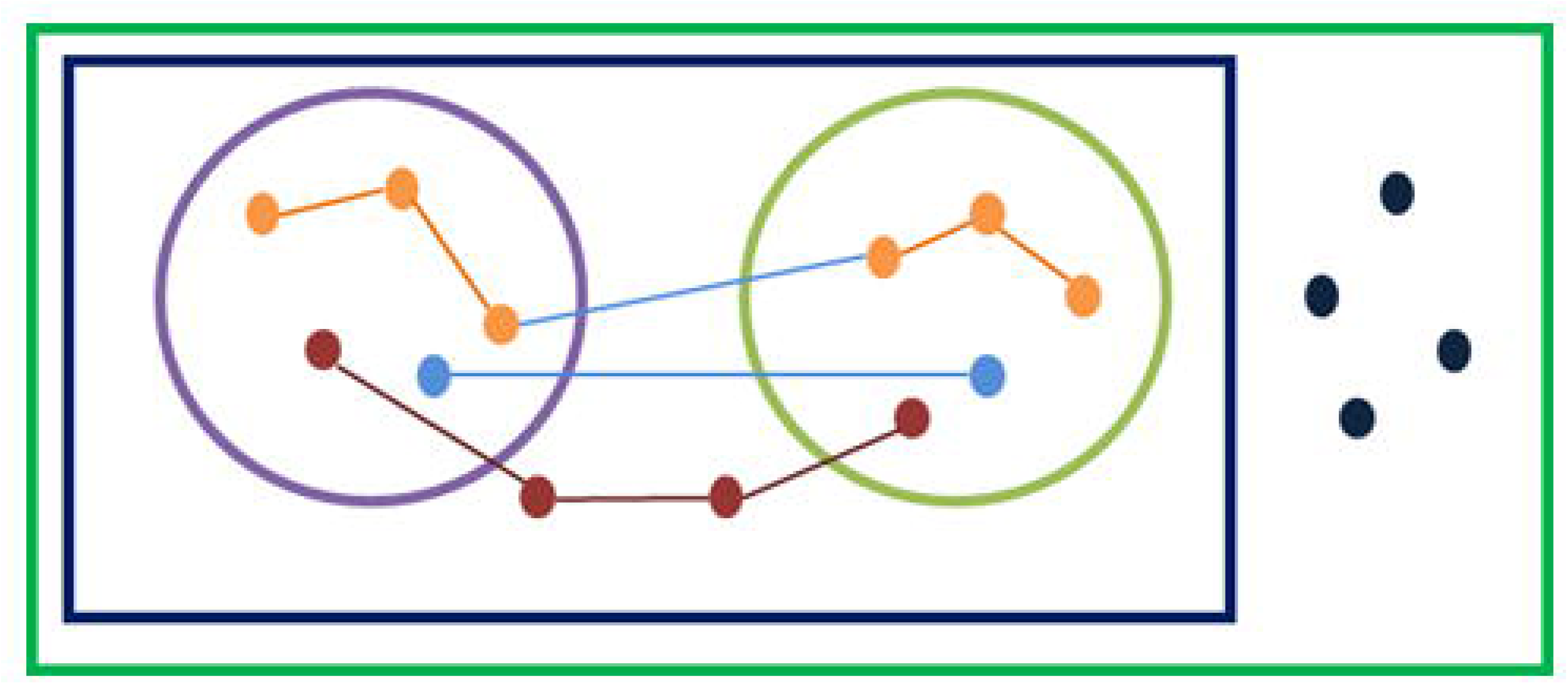
Schematic defining sets of genes and gene-pairs. Outermost rectangle represents all genes. Inner rectangle represents all genes that are in spatial proximity to at least one other gene. Circles represent annotated pathways. Nodes represent genes and edges represent spatially proximal gene pairs. All spatially proximal gene pairs are partitioned into three groups: ***proximal-intra-pathway:*** intra-pathway spatially proximal gene pairs (orange nodes and edges), ***proximal-inter-pathway:*** Spatially proximal pair of genes where each gene is in a different pathway (light blue nodes and edges), and ***proximal-generic:*** Spatially proximal pair of genes such that at least one of the genes is not in any pathway (red nodes and edges). Similarly all non-proximal gene pairs are partitioned into two categories: ***non-proximal-intra-pathway:*** a non-proximal gene pair within a pathway, and ***non-proximal-generic:*** any pair of genes that are not spatially proximal to any other gene and not within any pathway (pairs of dark blue nodes).

**Figure 6.**
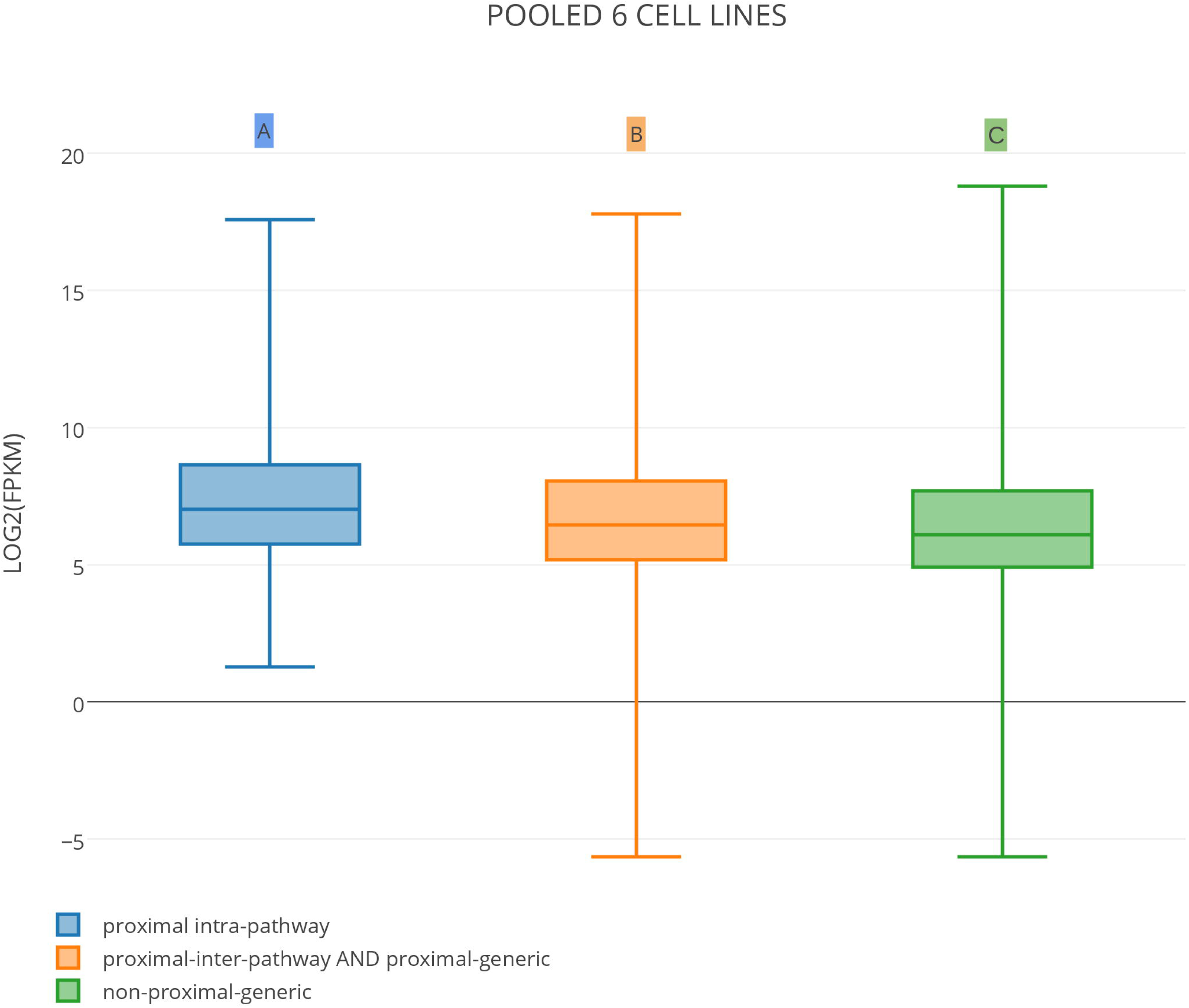
Spatial proximity and gene expression. The figure shows box-plots of gene expression (FPKM) values of the genes in three different groups (see Figure-5) pooled from all 6 cell lines. A∼B Wilcoxon test p-values = 3.4E-35. A∼C Wilcoxon test p-value = 8.43E-72. See Supplementary Fig. 4 for results of 6 individual cells.

We directly assessed the correlation between cell type specific pathway proximity and pathway activity. We used two measures to approximate pathway activity (see Materials & Methods). The first measure captured the mean expression of all genes in the pathway and the second measure captured the ratio of mean expressions of proximal and non-proximal genes in the pathway. Each measure was converted into a based on random sampling (see Materials & Methods). As shown in Fig. 7, spatial proximity is highly correlated with mean pathway expression (Spearman rho = 0.77, p = 5.9e-62). Note that this analysis does not rely on any Z-score cutoff.

**Figure 7.**
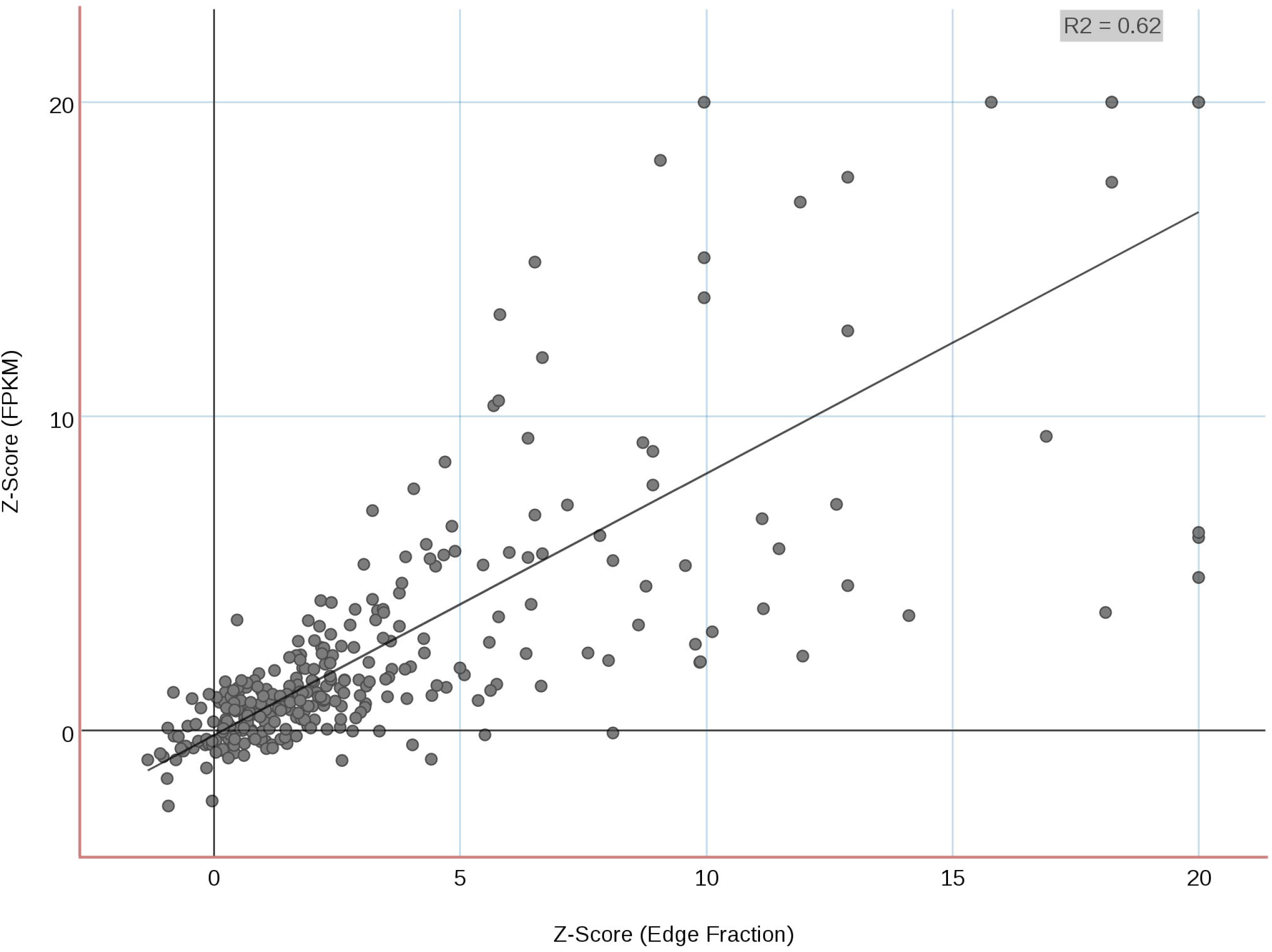
Spatial proximity versus mean pathway expression. This figure shows scatter plot between proximity z-scores of pathways versus z-scores of expression values of the proximal-intra-pathway genes pooled from all 6 cells. Spearman rho = 0.77, p-value: 5.90e-62.

Our observed highly positive correlation between spatial proximity of a pathway genes and the pathway activity would imply that the pathway genes are localized in active compartments and not in repressive or inactive compartments (in which case spatial proximity would result in pathway suppression – an opposite effect). A previous paper [48] has shown that in GM06990, relative to the inactive (B) compartment, the active (A) compartment (1) is highly enriched for genes, (2) has higher expression, (3) has more accessible chromatin, and (4) is loosely packed, i.e., has fewer interactions. To assess our hypothesis that interacting pathway genes are preferentially in A-compartment, we estimated the compartmentalization in GM06990 cell line using HOMER tool and found 18256 (75%) genes in A-compartment and 6371 (25%) genes in B-compartment, consistent with previous report. Next, we compared for 4 groups of genes (defined in Fig. 5) the tendency to belong to A-compartment. As shown in Supplementary Fig. 4 there is a robust monotonic trend whereby the genes proximal to another gene in the same pathway have the greatest tendency to belong to A-compartment and both spatial proximity and co-pathway membership contribute to this tendency. Specifically there a large difference between the first class (proximal-intra-pathway) and the non-proximal genes (p-value = 1.6E-53, Odds-ratio = 6.3). Thus, it seems that our observed positive correlation between pathway proximity and activity is due to the fact that most within-pathway proximal genes are in A-compartment.

### 3. Spatial proximity and protein-protein interaction

We have found that genes in a pathway tend to be spatially proximal. Previous studies have shown that the proteins in a pathway have a greater tendency to physically interact with each other [30]. We therefore directly assessed the correlation between spatial proximity of a gene pair and the physical interaction of their products. We obtained the protein-protein interactions (PPI) from HPRD database [31] and STRING [32]. Fig. 8 shows, for pooled data from all the 6 cell lines, for each of the 5 groupings of gene pairs (defined in Fig. 5), the fraction of all gene pairs that have evidence for physical interaction (fractional PPI). Results for other individual cell lines are provided in Supplementary Fig. 5. Taken together, these data suggest that, both pathway membership and spatial proximity of a gene pair is equally associated with PPI between their products, and the effect is partly independent, that is, PPI tendency is much greater for gene pairs that are both in the same pathway and physically proximal.

**Figure 8.**
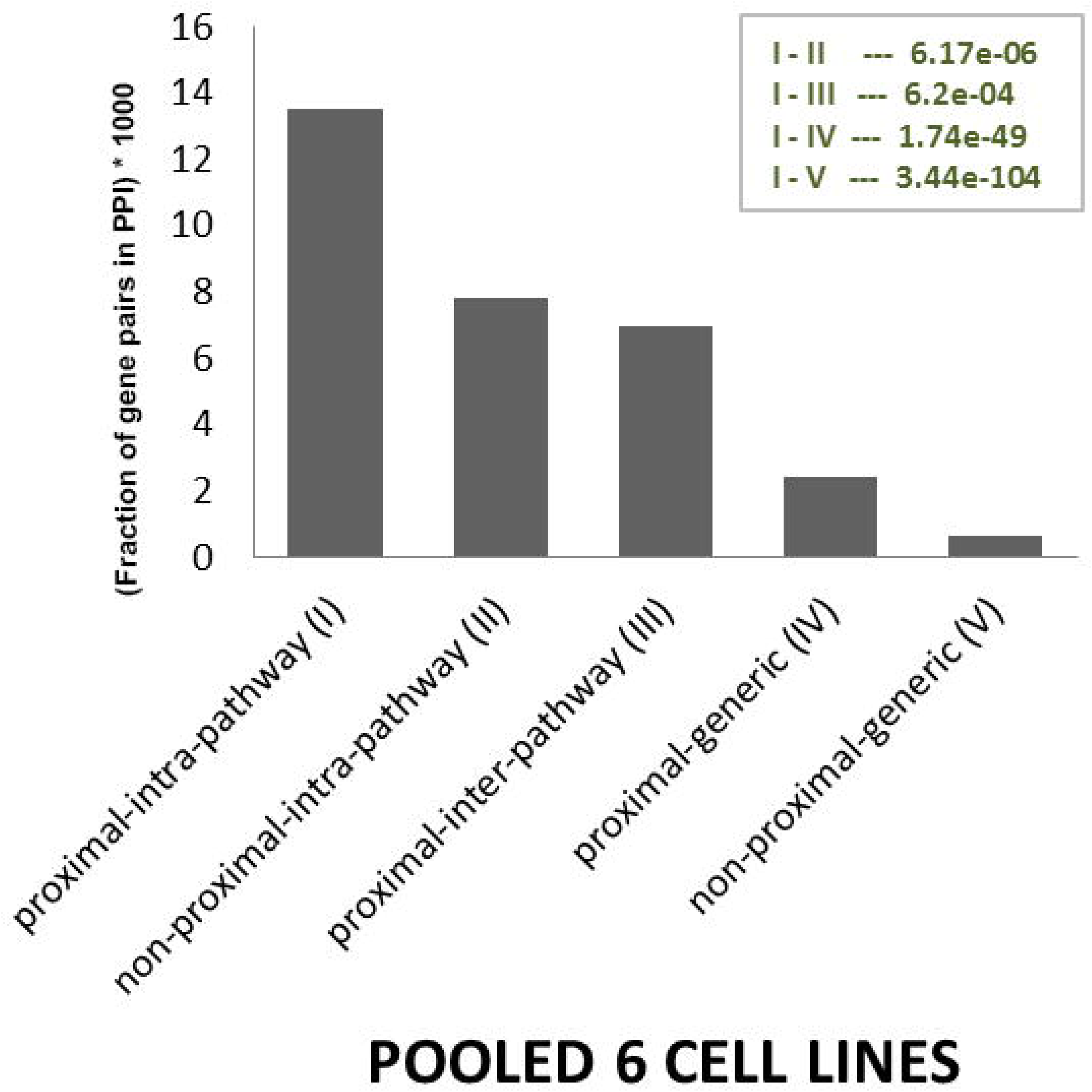
Pathway membership, spatial proximity and PPI. The figure shows 6 cells pooled fraction of gene pairs (Y-axis) in different gene groups (X-axis; see Figure 5) whose protein products physically interact. The inset shows Fishers tests p-values for various pair-wise comparisons. See supplementary Figure 6 for other tissues.

### 4. Functional enrichment for spatially proximal genes in pathways

Only a small fraction of genes in a pathway are spatially proximal to other genes in the same pathway. This could partly be due to low coverage and false negatives in the Hi-C derived interactions. However, given the differences between proximal and non-proximal genes above, we investigated whether specific functional terms are enriched among *proximal-intra-pathway* genes relative to non-proximal-intra-pathway genes (see Fig. 5 for definition). We pooled the data for all pathways into two groups and performed the functional enrichment analysis using GOrilla software [33]. Fig. 9 suggests that regulatory (both RNA processing, and splicing) and protein binding functions are highly enriched among the spatially proximal pathway genes. We emphasize that our background set of genes in this analysis included those that are in the pathways and are spatially proximal to some other gene not belonging to the same pathway. Thus the observed functional enrichment is not due to spatial proximity alone and is a specific property of spatially proximal genes within pathways.

**Figure 9.**
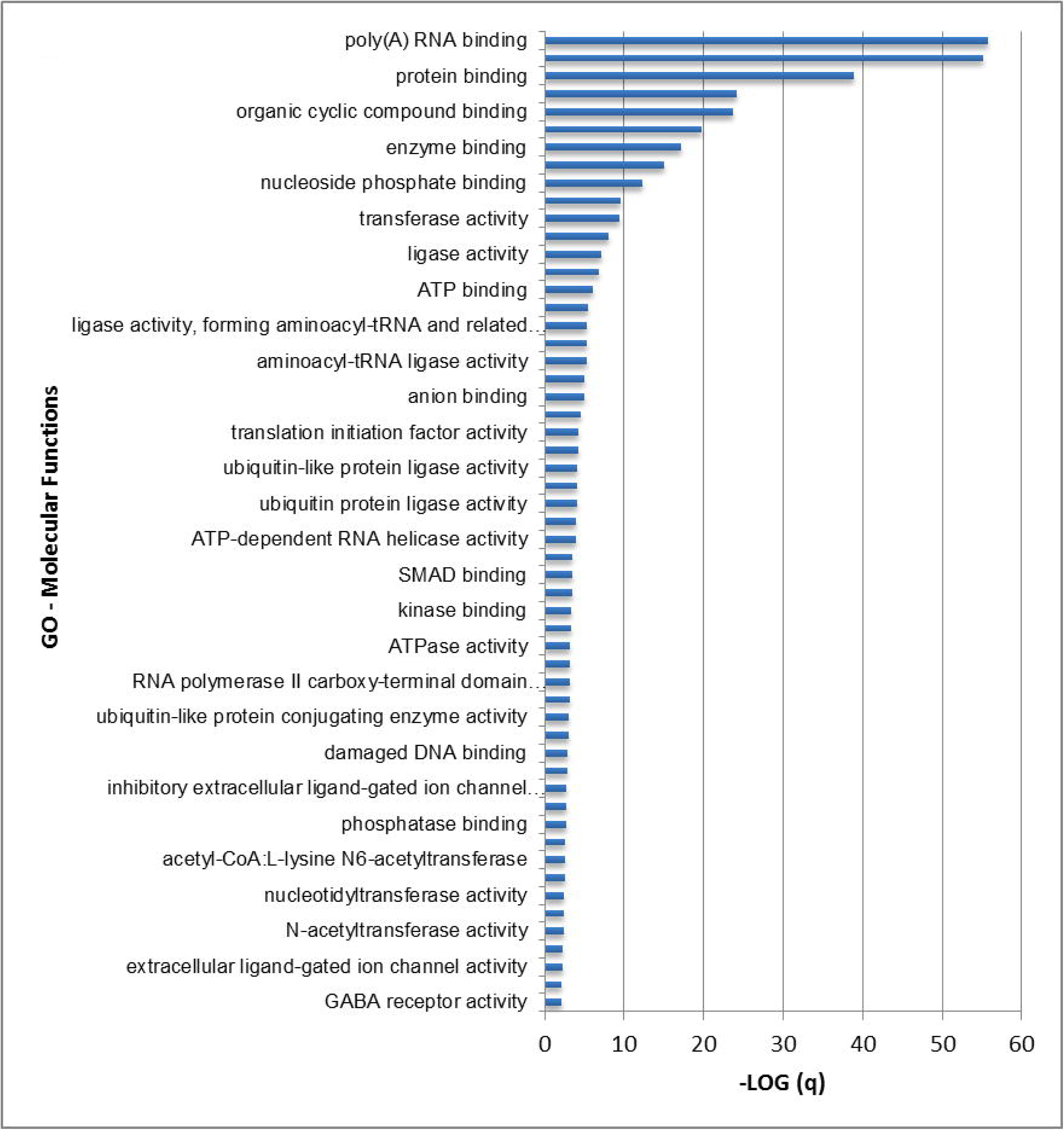
GO enrichment analysis. This figure enriched GO terms (–log (q-value) >= 2) ***proximal-intra-pathway*** genes relative to other spatially proximal genes (see Fig. 5).

### 5. Spatial proximity and regulatory hierarchy

The comparative analyses of biological properties of spatially proximal pathway genes relative to other pathway genes thus far suggests that the upstream genes in a pathway may be more likely to be spatially proximal, ensuring their robust expression and consequently robust pathway activity. We derived the hierarchical level of all pathway genes based on directed pathway edges (see Materials & Methods), and compared the hierarchical levels of proximal and non-proximal genes, pooled overall pathways. Our approach for assigning hierarchical level does not partition the pathway into a strict hierarchy, and can accommodate cycles. We found that, in 3 of the 6 cell lines, the hierarchical levels of proximal genes were higher than the rest; Wilcoxon test p-values: IMR90 (p = 4.4E-04), GM06990 (p = 9.7E-04) and hESC (p = 0.05). Lack of significance in other cell lines (p-values = 0.07, 0.16, and 0.2) may be attributed to due to insufficient data; when we pool the data from the other 3 cell lines, the result is significant (p = 1.41E-05). If we apply Chi-square test based on hierarchy level of 2 as the partitioning criterion, an additional cell line yields significance and importantly the odds ratio in all six cell lines range favorably from 1.7 to 3.2.

### 6. Spatial proximity between pathways

Next, we investigated, whether certain pairs of pathways might occupy neighboring spaces in the nucleus, suggesting their functional relatedness. Analogous to intra-pathway spatial proximity estimation, for each pair of pathways, after excluding the common genes, we obtained the number of interactions between genes across pathways and estimated its Z-score based on 1000 controlled random samplings (for computational tractability) of 2 sets of genes representing the 2 pathways. Using Z-score > 2 as the threshold, we identified a total of 3109 pathway pairs across all cell lines, 73 of which were proximal in at least 4 tissues and 20 were proximal in at least 5 tissues (Supplementary Table 2).

Next, analogous to intra-pathway proximity analysis, we estimated inter-pathway spatial proximity between housekeeping genes and all other KEGG pathways. We found that housekeeping genes are significantly proximal to 95 out of 164 KEGG pathways, at Z-score > 2 threshold in at least 1 cell line; the full distribution of number of housekeeping-proximal pathways shared in 1 or more cell lines is provided in Supplementary Fig. 6, and pathways proximal to housekeeping genes in 4 or more cell lines are provided in Supplementary Table 2. These results suggest a central role housekeeping genes may play in the genome organization as well as in coordinating the activities of other pathways.

Several of the pathway pairs deemed to be spatially proximal in our analysis (Supplementary Table 2) are very likely to be functionally related, for instance, pathways for metabolism of various amino acids and derivatives thereof. Importantly, however, these data also reveal non-trivial relationships, which have some support in literature. We discuss a few next.

- Consistent with the proximity of steroid hormone metabolism and proteolysis, direct functional links between these two processes have been noted [34].
- Links between amino acid metabolism and cell cycle and cancer (Chronic myloid leukemia in our case) is not surprising given the metabolic requirements during cell division and growth.
- JAK-STAT pathway and SNARE complexes are found to be proximal. While we did not find a direct link between the two, they are known to be co-targeted by TGF-beta signaling [35].
- Abnormal *steroid hormone metabolism* has been reported in *Myeloid leukemia* patients [36], which is consistent with the detected proximity of these two pathways.
- *Butanoate metabolism and Glioma* are deemed to be spatially proximal in our analysis. Treatment with butanoate (sodium butyrate) is known to induce differentiation of c6 Glioma cells [37].
- While a direct link between *Cholera* and *DRPLA*, deemed to be proximal, is not clear, we note that *DRPLA* is a spinocerebral degeneration disease, and *cholera toxin subunit b* is known to be dispersed to brain and spinal cord neurons [38].
- *Synthesis and degradation of ketone bodies* was found to be proximal to genes involved in several different cancers, including pancreatic cancer, and cell cycle. Previous papers have shown growth inhibitory effects of *ketone* bodies on pancreatic cancer could be mediated by reduced c-Myc expression [39]. Moreover c-Myc is overexpressed in pancreatic cancer [40] and its inhibition has been shown to result in regression of *lung cancer* [41] and pancreatic cancer [42].
- *Glycosylphosphatidylinositol (GPI)* and *Huntington* (a neurological disease) were deemed proximal. While a direct link between these two pathways is not clear, GPI-anchor cleavage is known to modulate notch signaling and promoter neurogenesis [43].

Overall, these results suggest that spatial proximity between pathway genes is somewhat associated with functional interactions between the pathways.

## Discussion

In this work, we have presented the first comprehensive analysis of intra- and inter-pathway spatial proximity in multiple *Homo sapiens* cell lines. Previous studies have shown that in *Saccharomyces cerevisiae* [10], *Plasmodium falciparum* [12] and *H.sapiens* lymphoblastoid cell line [13], broadly, functionally related genes tend to be spatially proximal. Our goal here was to not only extend these previous observations to multiple human cell lines and assess the relationships between spatial proximity and pathway activity based on gene expression, but equally importantly, to further functionally characterize proximal genes within pathways and examine higher-order physical and functional interactions between pathways.

Previous similar analysis in *S. cerevisiae* [10] are based on only inter-chromosomal segment interactions, and *H. sapiens* lymphoblastoid cell line results [13] are based on low-resolution (1 Mb) segments, which can result in spurious interactions at the level of individual gene loci. Importantly, however, a greater tendency for genomically proximal regions to be spatially proximal, i.e., autocorrelation, unless appropriately controlled for, can result in false positives in inferring significant spatial proximity from Hi-C data. The absence of effective tools to control for autocorrelation has forced previous studies to exclude intra-chromosomal interactions from consideration, significantly impacting their statistical power [10]. In contrast the Homer tool [20] satisfactorily controls of autocorrelation in estimating significance, consequently enabling us to include all significant interactions, both inter- and intra-chromosomal, in our analyses, while obviating an explicit control for inter-gene distances in random sampling procedures. We note that despite an explicit control for autocorrelation, genomically proximal regions (within 500 kb) have a slightly higher tendency to be spatially proximal (data not shown). However, we explicitly tested if this biases our pathway proximity assessment as follows. We compared intra-pathway gene distances with those for randomly selected genes from the same chromosome and found that for none of the pathways there was a significant difference between the two sets of distances.

We have performed our analyses based on 100 kb resolution to detect interactions. However, we have also assessed the impact of using a higher resolution of 10 kb. We found that the number of interactions detected was much greater when using 100 kb resolution, especially the inter-chromosomal interactions, due to a greater statistical power for interaction detection at this resolution. For example, for cell type HEK293, only 7889 intra- and only 14 inter-chromosomal gene-gene interactions are detected using 10kb resolution and 43439 intra- and 1538 inter-chromosomal interactions at 100 kb. Moreover, at 10 kb resolution, large fractions of detected interactions are within a gene, which does not contribute to our analyses. We note that using a 100 kb resolution does not substantively influence the gene-gene interaction inference, as only 4% of 100 kb segments have multiple genes. Thus, to maximize statistical power with relative small fraction of ambiguous (but not necessarily biologically wrong) gene-gene interaction calls, we chose to perform all downstream analyses based on 100 kb resolution.

Hi-C data, like most genome-wide datasets, comes with a level of false positives, as does the pathway data. This is an important issue, and one that cannot be addressed by computational means alone. We have relied on the published interaction data that have undergone quality control measures, and used robust tool with recommended controls to perform the analyses. For pathway proximity, we have relied on a well-controlled randomized gene set to assess the significance of proximity. Despite the controls, a certain fraction of data is likely to be false positive. However, the noise in the data, as long as the tests are properly controlled, is not likely to generate strong consistent signals across multiple cell types simply by chance.

We performed a number of checks to ensure the robustness of our conclusions against several potential biases. First, note that in quantifying the significance of the pathway spatial proximity, we control for the lengths of the pathway genes. Second, we ensured that our ability to detect the spatial proximity of a pathway is independent of the number of genes in the pathway (Spearman correlation between pathway size and spatial proximity Z-score was statistically insignificant). A recent paper [9] has suggested a link between detection of Hi-C interaction of a gene and a gene’s codon usage (which is related to its expression) and the GC content of the gene’s genomic locus. Third, we ascertained that the codon-usage does not bias the detection of Hi-C interactions (Spearman correlation = -0.09).

Finally, with regards to the potential GC bias, indeed highly interacting loci tend to have lower GC composition, as noted previously [11]. Although the GC composition near the restriction sites can present a technical bias [44], several papers also suggest that GC composition may be an inherent property of the physical proximity [45]. Therefore, it seems that an explicit control for GC content may not be ideal. However to explore the extent to which GC-controlled analyses would affect our results, for HEK293 cell line we reprocessed the data with specific GC control option provided in HOMER tool and compared the downstream results of our analyses with and without the GC control. We found that 80% of interactions detected without GC control are also detected with GC control and overall correlation between gene-wise interaction degrees between the two is 0.92. Next, we re-estimated the pathway proximity Z-scores based on GC-controlled interactions and found those to be highly correlated with the Z-score based on uncontrolled interaction detection (Spearman rho = 0.85). Lastly, we recalculated the correlation between pathway proximity and pathway activity and found that too is as significant as the estimated without GC control (Spearman rho = 0.80, relative to 0.83 without GC control). Thus, our conclusions are not substantively biased by these various potentially confounding factors.

Our primary resource of biological pathways – KEGG, is dominated by essential and broadly utilized cellular pathways, and therefore it is encouraging to see that by and large KEGG pathways are not only highly significantly proximal (Fig. 2), but a large fraction of these pathways are proximal in multiple cell lines (Fig. 4). However, as we show, this is not true for a different set of pathways relevant to cancer, where the overall z-scores are much more subdued, and spatially proximity is less ubiquitous (Supplementary Fig. 2). Despite, general ubiquity of spatial proximity of KEGG pathways, we still see a strong correlation between cell type-specific spatial proximity and pathway activity as approximated by gene expression (Fig. 7).

Previous studies have noted greater transcription in spatially clustered regions [8]. Independently, earlier studies have shown the existence of so called transcription factories [46] - nuclear locales with enriched core transcriptional machinery components where transcripts are synthesized and processed. Moreover, links between transcription factories and chromatin organization have been noted [47]. Taken together, these previous results are consistent with our observation that genes in spatially proximity with other genes have much higher expression. However, interestingly, in addition to spatial proximity alone, functional relationship between the spatially proximal genes, i.e., membership in the same pathway, makes a small but significant additional contribution to the gene expression level (Fig. 6).

Our analysis reveals an unexpected association between spatial proximity of a gene pair and the interaction between their protein products. Among the physically proximal gene pairs, the well-annotated genes, i.e., annotated in some KEGG pathway, have much greater PPI propensity than genes that do not belong to an annotated pathway ((III) vs. (IV) in Fig. 8); this may be explained by a greater representation of well-studied genes in PPI databases. Functionally related genes have been shown to have a greater propensity to physically interact [49], consistent with our findings (compare (I) and (III) in Fig. 8). However, we found that spatial proximity is independently associated with protein interaction in both pathway ((I) versus (II) in Fig. 8) and non-pathway ((IV) versus (V) in Fig. 8) contexts. The gene-pairs that are both spatially proximal and belong to same pathway have the highest PPI propensity ((I) in Fig. 8). These trends are identical in all 6 cell lines, suggesting that both spatial proximity and pathway membership contribute independently to PPI propensity.

Scrutinizing each of the pathways, we found the spatially proximal genes in a pathway to have distinguishing functional characteristics relative to other pathway genes that are not spatially proximal to any other gene in the same pathway. In terms of biological processes (Fig. 9), such genes are overwhelmingly involved in transcription, splicing and intracellular transport and localization. Interestingly, the genes in a pathway that are spatially proximal to other genes in the same pathway tend to occupy a higher level in the regulatory hierarchy, related to other genes in the pathway, that also are spatially proximal to other genes but none in the same pathway. Overall, these results ascribe, for the first time, a special functional status to spatially proximal genes in pathways – such genes tend to perform higher-level regulatory functions. We found that housekeeping genes, consistent with their ubiquitous expression and activity, tend to be broadly and highly spatially proximal. Previous studies have observed clustering of housekeeping genes into so call transcription factories [50], and have suggested that interactions between housekeeping genes may play a role in the spatial organization of the chromatin [51]. Our results confirm these previous observations through the first genome-wide assessment of spatial proximity of housekeeping genes in multiple cell lines. In addition to spatial proximity of housekeeping genes, we also found that as a group housekeeping genes are ‘centrally’ located in the nucleus and act as a link between numerous other pathways. A mechanistic interpretation of this intriguing observation, as well as of the causal link between the expression of housekeeping genes and their spatial proximity, will require further analysis.

## Conclusion

Overall, based on the first comprehensive pathway-centric analysis of spatial proximity in multiple cell lines, our results suggest that (i) context-specific regulation of pathways is associated with their context-specific spatial proximity; in doing so, our analysis provides mechanistic insights into cell type-specific activity of certain pathways, (i) spatial-proximity of pathway genes is associated with physical interaction among their gene products, and (iii) specific classes of genes within pathways, likely occupying higher regulatory levels, have a greater tendency to be spatially proximal. Our results also provide insights into correlated activity of multiple pathways by showing that the genes in these pathways are spatially proximal.

## Methods

### Hi-C processing pipeline

We downloaded paired-end Hi-C raw reads FASTQ files of sample replicates for the following tissues from GEO database (www.ncbi.nlm.nih.gov/geo): (i) HEK293 (GSM1081530, GSM1081531) [15], (ii) IMR90 (GSM1055800, GSM1055801) [52], (iii) hESC (GSM862723, GSM892306) [4], (iv) GMO6990 (GSM1340639) [16], (v) RWPE1 (GSM927076) [17] and (vi) BT483 (GSM1340638, GSM1340637) [16]. We mapped the reads onto hg19 human genome using *BWA* tools [53] with default parameters. The resulting SAM files were converted to BAM files using “*samtools view*” program [54] and processed to removing PCR duplicates using “*samtools sort*” and “*Picard*” tools.

We then processed the non-redundant reads for Hi-C analysis using various HOMER tools [20]: we ran “*makeTagDirectory*” program using options “*tbp -1*” (to ensure that any genomic location is mapped by a unique read), *“–restrictionSite*” (only keep reads if both ends of the paired-end read have a restriction site within the fragment length estimate 3’ to the read), “*-removePEbg*” (removing read pairs separated by less than 1.5x the sequencing insert fragment length), “-removeSelfLigation” (remove re-ligation events), and “*-removeSpikes*” (remove high tag density regions).

### Normalization of Hi-C interactions

Having output from the previous steps, next we normalized the data to create background of the Hi-C interactions at 100kb resolution using HOMER “*analyzeHiC*” program. The program (i) divides the genome into 100kb regions, (ii) calculates total read coverage in each region, (iii) calculates the fraction of interactions spanning any given distance with respect to read depth, (iv) optimizes a read count model to assign expected interaction counts in regions with uneven sequencing depth and (v) calculates variation in interaction frequencies as a function of distance. For fragments i and j, the procedure the estimated expected number of reads supporting the interaction as:

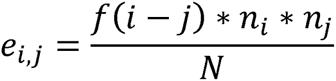

where *N* = total number of reads, *n* = number of reads in a region, and *f* represents the expected frequency of Hi-C reads as a function of distance.

The background model were created for the entire data for a given sample then applied for selection of Hi-C interacting reads at default p-value cutoff of 0.001 using “*-interaction*” parameters using “*analyzeHiC*” program. We further filtered the Hi-C interactions using FDR cutoff <= 0.1 and considered the resulting set as significant Hi-C interactions. The significant Hi-C interacting reads were then passed through “*annotateInteractions*” program for mapping them onto the annotated genomic features (i.e., 5’UTR, CDS, introns, exons, intergenic etc.), from which, we selected only those interactions mapping to gene regions (i.e., Promoter, 5’UTR, exon, intron and 3’UTR), resulting in a comprehensive set of spatially proximal gene pairs (SGP) for each tissue.

### Codon usage

To assess the effect of codon bias on interaction detection, we used Biopython’s “*SeqeUtils::ModuleCodonUsage*” package and calculated Codon Adaptation Index (CAI) for each gene of the H. sapiens genome.

### Pathway datasets

A ‘pathway’ for our purpose is a set of genes. We downloaded two sets of pathways from KEGG [21] and NetPath [19] databases, only retaining those with at least 10 genes, resulting in 164 and 32 pathways respectively. We also included the set of 3800 housekeeping genes [22], as an additional ‘pathway’.

### Defining classes of edges and nodes relative to pathways

For various analyses we have defined 5 disjoint sets of genes and gene pairs in the context of a pathway and spatial proximity (see Figure 5 for illustration). All spatially proximal gene pairs were partitioned into three groups: *proximal-intra-pathway:* intra-pathway spatially proximal gene pairs, *proximal-inter-pathway:* Spatially proximal pair of genes where each gene is in a different pathway, and proximal-generic: Spatially proximal pair of genes such that at least one of the genes is not in any pathway. Similarly all non-proximal gene pairs were partitioned into two categories: *non-proximal-intra-pathway*: a non-proximal gene pair within a pathway, and *non-proximal-generic*: any pair of genes that are not spatially proximal to any other gene and not within any pathway.

### Edge fraction and its significance in estimating pathway spatial proximity

Intra-pathway gene spatial proximity was estimated as *Edge Fraction* (EF) – number of pairwise gene-gene interaction in the pathway normalized by the number of total possible interactions. To quantify significance of the *EF* for pathway with *N* genes, we randomly sampled *N* genes, such that number of genes in each chromosome is identical to real pathway, and each sampled gene’s length was within 20% of the matched pathway gene’s length (this controls for length-based bias in interaction detection). We generated 1000 such samples and calculated 1000 corresponding *EFs*. We then obtained the Z-score corresponding to the EF for actual pathway relative to 1000 controls. We estimated the Z-score for each pathway in each cell line resulting in 165 × 6 Z-score matrix.

### Inter-pathway proximity

Analogous to the intra-pathway proximity analysis above, for a pair of pathways, after excluding the shared genes, we estimate the edges between genes in two pathways, and estimate its significance based on randomly sampling 2 gene sets (instead of 1 as above), with identical controls as above. We thus estimated a Z-score for inter-pathway proximity for all pairs of pathways.

### Processing RNA-Seq data

We downloaded raw RNA-Seq FASTQ files for three of the tissues in which matching RNA-Seq was available: (i) HEK293 (GSM1081534, GSM1081535) [15], (ii) IMR90 (GSM1154029) [52] and (iii) RWPE1 (GSM927074) [17], (iv) hESC (GSM758566) [55], (v) GM06990 (GSM958747) [56] and (vi) BT483 (GSM1172854) [57]. The raw FASTQ reads were mapped and processed up to de-duplication steps using the same pipeline that we used to apply for Hi-C data analysis, and then used to quantify expression levels using cufflinks [58] tool with default parameters, yielding gene-wise RPKM values.

### Significance of pathway activity

For genes with multiple transcripts, we take the maximum expression over all transcripts for the genes. For a pathway, we estimated its activity as the average expression of the genes in the pathway; we selected only the *proximal-intra-pathway* genes to estimate activity. To quantify significance of the pathway activity, we followed a sampling approach similar to that for estimating EF significance above, yielding a z-score for each pathway and cell line.

### Estimating regulatory hierarchy

The regulatory analysis of a pathway requires a directed graph in which all defined interactions suggest direction of the signal flow. KEGG does not provide the directed graph by default. Therefore, we downloaded *KGML* (KEGG Markup Language) and parsed the files using *KEGGgraph* package in Bioconductor and created directed graph for each pathway using *NetworkX* package in Python. In order to investigate hierarchy of genes we first assigned a synthetic root connecting to all pathway nodes with zero in-degree, and then calculated the shortest path length (SPL) from root to every other node. Low SPL indicates higher level of hierarchy. For Chi-square and Fisher tests (see Results), we created two gene sets: (i) genes with SPL = 1 (top level) and (ii) genes with SPL >= 3 (lower hierarchy). We pooled these two sets across all pathways.

## Author contributions

S.H. conceived the project with input from C.K. and M.G. H.K. designed and implemented the software pipeline and performed all analyses. S.H. wrote the manuscript with help from all authors.

## Additional Data Files

“Supplementary File” contains supplementary figures and tables. File ‘Supplementary Fig1’ contains the Supplementary Fig. 1.

## Acknowledgement

This work was funded by NIH HG007104 to C.K. and NIH GM100335 to S.H.

